# Longitudinal assessment of sputum microbiome by sequencing of the 16S rRNA gene in non-CF bronchiectasis patients

**DOI:** 10.1101/050237

**Authors:** Michael J Cox, Elena M Turek, Catherine Hennessy, Ghazala K Mirza, Phillip L James, Meg Coleman, Andrew Jones, Robert Wilson, Diana Bilton, William O. C. Cookson, Miriam F. Moffatt, Michael Loebinger

**Affiliations:** National Heart and Lung Institute, Imperial College London, UK, SW3 6LY; Royal Brompton and Harefield NHS Foundation Trust, London, UK

**Keywords:** 16S rRNA gene sequencing, respiratory, chronic suppurative lung disease, bronchiectasis, non-cystic fibrosis bronchiectasis, longitudinal

## Abstract

**Background:** Bronchiectasis is accompanied by chronic bronchial infection that may drive disease progression. However, the evidence base for antibiotic therapy is limited. DNA based methods offer better identification and quantification of microbial constituents of sputum than standard clinical culture and may help inform patient management strategies. Our study objective was to determine the longitudinal variability of the non-CF bronchiectasis microbiome in sputum with respect to clinical variables.

Eighty-five patients with non-cystic fibrosis (CF) bronchiectasis and daily sputum production were recruited from outpatient clinics and followed for six months. Monthly sputum samples and clinical measurements were taken, together with additional samples during exacerbations. 16S rRNA gene sequencing of the sputum microbiota was successful for 381 samples from 76 patients and analysed in conjunction with clinical data.

**Results:** Microbial communities were highly individual in composition and stability, usually with limited diversity and often containing multiple pathogens. When compared to DNA sequencing, microbial culture had restricted sensitivity in identifying common pathogens. With some exceptions, community characteristics showed poor correlations with clinical features including underlying disease, antibiotic use and exacerbations, with the subject showing the strongest association with community structure. When present, certain pathogens may also shape the structure of the rest of the microbial community.

**Conclusions:** The use of microbial community analysis of sputum added to information from microbial culture. A simple model of exacerbations driven by bacterial overgrowth was not supported, suggesting a need for revision of principles for antibiotic therapy. In individual patients, the management of chronic bronchial infection may be improved by therapy specific to their microbiome, taking into account pathogen load, community stability, and acute and chronic community responses to antibiotics.

## Background

Bronchiectasis is characterised by abnormal dilated thick-walled bronchi and is often accompanied by chronic bronchial infection. Patients with advanced disease may produce copious volumes of purulent sputum and lung function may be severely and progressively impaired. The prevalence of non-CF bronchiectasis in the US has been estimated at 272 per 100 000 persons over 75 years of age and hospitalisation rates are increasing (Seitz 2010).

Although chronic infection with episodes of exacerbation may drive the progression of bronchiectasis, the evidence base for antibiotic therapy is limited(Pasteur et al. 2010). An underlying assumption is often that exacerbations are driven by the overgrowth of a particular microbial species, although mixed pathogen colonisations are recognised. Antibiotic choice in current practice is initially empirical until sputum cultures are obtained and then directed by isolated organism (Pasteur et al. 2010).

Standard microbial cultures are selective, identifying a restricted range of bacterial species in clinical samples. Molecular, culture-independent, techniques such as 16S rRNA gene sequencing have been shown to detect a much greater variety of microbes from the same specimens as standard culture techniques (Huang et al. 2010; Erb-Downward et al. 2011; Tunney et al. 2011; Cox et al. 2010; Charlson et al. 2010; Hilty et al. 2010; Rogers et al. 2009).

In order to understand the potential impact of culture-independent techniques on the management of chronic bronchial infection, we have carried out a prospective six-month study of patients with CT-defined bronchiectasis attending clinics at the Royal Brompton Hospital, London. Recruited patients were studied at monthly intervals and during any exacerbations.

We present here the results of quantification of the bacterial burden by quantitative PCR of the 16S rRNA gene and community analyses, comparing them to clinical outcomes and microbiological cultures.

Microbial diversity reflects the number of species, their presence and their abundance, in a study. Higher levels of diversity are associated with resilience of microbial communities to invasion, and may characterise human health. We have therefore examined associations with the number and proportions of species in individual patients (captured by measures of α-diversity), and with the community structure across patients (reflected in (β-diversity statistics).

Some of this work has been presented before in the form of an abstract and presentation at the American Thoracic Society International Conference 2015 (Cox et al. 2015) and pre-print at BioArxiv.org (Cox et al. 2016)

## Methods

### Participants

We recruited patients with CT (computerised tomography) defined non-CF bronchiectasis and daily sputum production between December 2010 and May 2011 at the Royal Brompton Hospital, a tertiary referral centre. Patients gave their full informed consent and the study was approved by South West London REC under reference number 10/H0801/53. Patients had all previously been screened according to the British Thoracic Society bronchiectasis guidelines. Patients had monthly research visits at which fresh sputum samples were collected and spirometry and clinical assessment performed. Patients were encouraged to attend the centre for a suspected exacerbation and to provide a further sputum sample. An exacerbation was defined as an acute deterioration with worsening local symptoms (increased cough, sputum volume, viscosity, purulence, breathlessness) and/or systemic upset and the physician determined need for antibiotics as per the BTS bronchiectasis guidelines. The clinical state for each sample was defined as baseline (B), exacerbation (E), treatment (T), or recovery (R) as previously published(Zhao et al. 2012). Briefly, B was defined as well or mild increase in respiratory symptoms, no doctor defined respiratory exacerbation, not hospitalised, not on episodic antibiotics for more than 30 days. E was defined as a doctor defined respiratory exacerbation, the sample was prior to the start of episodic intravenous (IV) or oral antibiotics, not on episodic antibiotics for more than 30 days. T was defined as on IV or oral episodic antibiotics for treatment of doctor defined respiratory exacerbation whilst R was defined as off episodic antibiotics for less than 30 days and may or may not be back to baseline clinical state. Patients recorded antibiotic use during the six month period.

Sputum samples underwent the standard clinical microbiology and reporting for non-CF sputum samples (Chocolate agar and Blood agar at 5% CO_2_ and 37 ^o^C, MacConkey agar at 37°C). Samples for molecular testing were stored frozen at –80°C prior to DNA extraction which was performed using the FastDNA Spin Kit for Soil (MP Biomedicals, Santa Ana, CA, USA) as per manufacturer instructions (including 55°C incubation at elution). Bead-beating was performed at 6800 rpm for two cycles of 30 seconds (Precellys, Bertin Technologies, Montigny-le-Bretonneux, France).

### Molecular microbiology

Quantitative PCR of each extracted DNA was performed in triplicate using the Viia7 Real Time PCR system (Life Technologies, Waltham, MA, USA) and the primers 520 F AYT GGG YDT AAA GNG and 802 R TAC NVG GGT ATC TAA TCC targeting the 16S rRNA gene V4 region. Samples that failed to amplify were repeated twice to confirm the result. Reaction conditions are available in supplementary methods.

16S rRNA gene amplification of the V3–V5 region was performed in quadruplicate using adapted primers 357F/926R (Sim et al. 2012) with 12 bp barcodes included in the reverse primer (Fierer et al. 2008) and 454 sequencing adaptors A and B included in the reverse and forward primers respectively. Sequencing was performed on a Roche 454 (454 Life Sciences, Branford, CT, USA) and further methodology and sequence processing is detailed in supplementary methods. The resulting OTU table of 381 samples and 352 OTUs was used for all subsequent analyses and imported to the R statistical environment as a PhyloSeq (McMurdie & Holmes 2013) object, along with a mid-point rooted FastTree phylogenetic tree. A cross-sectional dataset of 72 samples and 194 OTUs was produced by sub-sampling the first baseline classified (B in the BETR scheme) sample available from each patient in PhyloSeq.

For comparisons with microbial culture, OTUs were selected by either being the most abundant OTU with similar identification (*Pseudomonas, Moraxella, Streptococcus*), the only OTU identified as that genus (*Stenotrophomonas, Proteus*) or the representative read positively identified by phylogenetic analyses as the same species (*Haemophilus influenzae*).

### Statistics

Statistical analysis was performed in the R statistical environment (version 3·1·2, (Team 2014)).

Species richness, Pielou’s evenness, Shannon’s Diversity Index, and Inverse Simpson’s Index were calculated for each sample. Diversity metrics were assessed for normality using Shapiro-Wilk’s tests and quantile-quantile plots. Shannon’s Diversity Index, Inverse Simpson’s Index and Pielou’s evenness were tested against variables using Wilcoxon Signed Rank and Kruskal Wallis Rank Sum tests. Species richness was normally distributed and tested using Student’s T-test and ANOVA. Spearman’s Rank correlations were used for continuous data. Bacterial load as determined by qPCR was normally distributed and paired Student’s T-test used; to do so, the first available consecutive samples of each combination of BETR class were used from each patient in this analysis i.e. B-B, B-E, B-T and B-R. If there was no available B sample in the month prior to another class, this pair was dropped.

For the Adonis (PERMANOVA) analyses, all variables were tested independently in the cross-sectional dataset. Those that proved to be significant were taken forward for further testing together in a larger model. Non-significant variables were backward removed from the models and samples with missing data were removed list wise. The order of the variables remaining was also tested as this can influence a PERMANOVA and the final model explains the maximum variance possible in the most limited number of variables. The final variables included in order were: isolation of mucoid *Pseudomonas aeruginosa,* isolation of *Haemophilus influenzae,* prophylactic treatment with Colistin and isolation of *Staphylococcus aureus* and the detailed model and R values can be found in the supplementary methods.

Sequence data has been submitted to the European Nucleotide Archive and is available under accession number PRJEB14304 and sample accessions ERS1201047 to ERS1201427. In addition the denoised reads, OTU table and clinical data are available to download at http://lungen.bioinformatics.ic.ac.uk/data/microbiomeLAMBbronchiectasis/

## Results

### Patient characteristics

Eighty-five subjects were recruited and produced 467 sputum samples. Post DNA and sequencing quality controls, 381 samples from 76 subjects were included in the longitudinal analysis. A cross-sectional dataset was created by taking the first baseline sample of 72 subjects. The remaining four subjects had only exacerbation, treatment or recovery samples. The aetiology of the bronchiectasis was most commonly idiopathic (47 %) or post-infective (25 %) (Table 1 and Figure 1). In general, the disease was severe (median baseline FEV1 % predicted 63 %, IQR 54 % to 82 %).

**Table 1.**
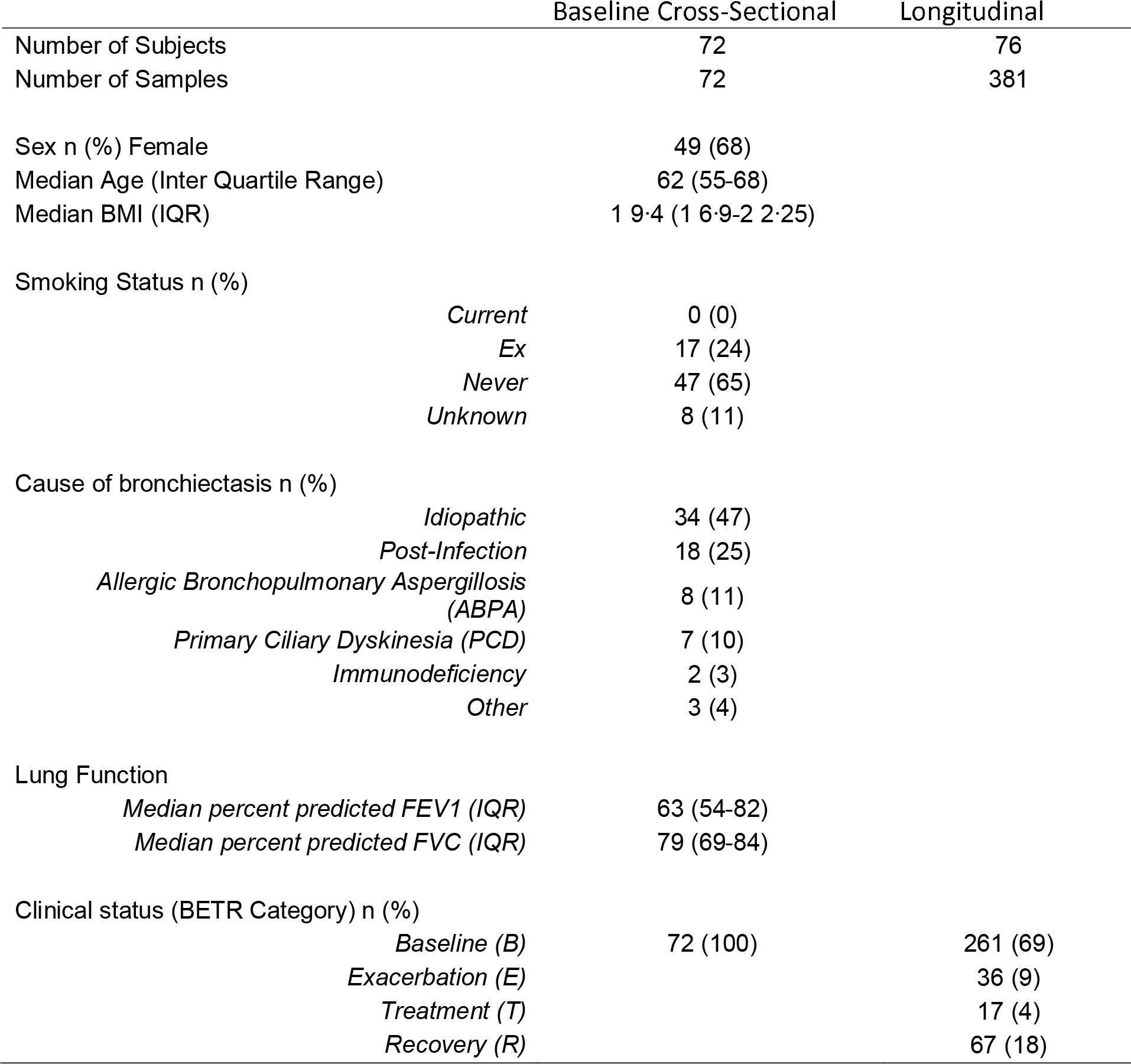
Patient Demographics

|  | Baseline Cross-Sectional | Longitudinal |
| --- | --- | --- |
| Number of Subjects | 72 | 76 |
| Number of Samples | 72 | 381 |
| Sex n (%) Female | 49 (68) |  |
| Median Age (Inter Quartile Range) | 62 (55-68) |  |
| Median BMI (IQR) | 19.4 (16.9-22.25) |  |
| Smoking Status n (%) |  |  |
| *Current* | 0 (0) |  |
| *Ex* | 17 (24) |  |
| *Never* | 47 (65) |  |
| *Unknown* | 8 (11) |  |
| Cause of bronchiectasis n (%) |  |  |
| *Idiopathic* | 34 (47) |  |
| *Post-Infection* | 18 (25) |  |
| *Allergic Bronchopulmonary Aspergillosis (ABPA)* | 8 (11) |  |
| *Primary Ciliary Dyskinesia (PCD)* | 7 (10) |  |
| *Immunodeficiency* | 2 (3) |  |
| *Other* | 3 (4) |  |
| Lung Function |  |  |
| *Median percent predicted FEV1 (IQR)* | 63 (54-82) |  |
| *Median percent predicted FVC (IQR)* | 79 (69-84) |  |
| Clinical status (BETR Category) n (%) |  |  |
| *Baseline (B)* | 72 (100) | 261 (69) |
| *Exacerbation (E)* |  | 36 (9) |
| *Treatment (T)* |  | 17 (4) |
| *Recovery (R)* |  | 67 (18) |

**Figure 1.**
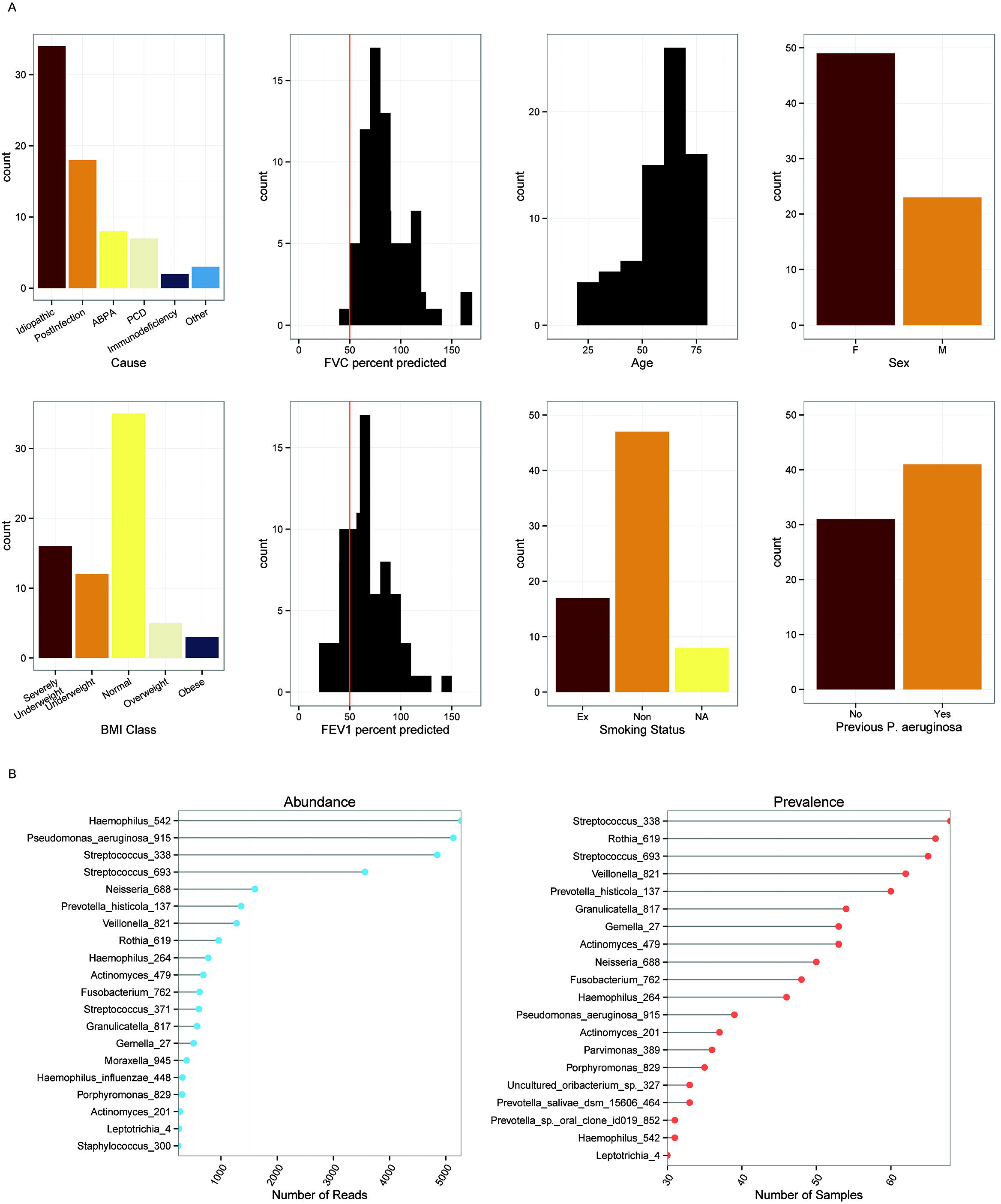
**IA.**Demographics of the non-CF bronchiectasis cohort indicating distribution of (from left to right, top to bottom): the cause of bronchiectasis; FVC percent predicted (red line indicates 50%); subject age; subject sex; BMI class; FEV1 percent predicted (red line indicates 50%); smoking status; and whether subject has previously cultured *P. aeruginosa*. **IB.**Distribution of OTUs within the cohort. Abundance is the total number of reads assigned to an OTU from any sample. Prevalence is how often an OTU is detected in samples. Haemophilus_542 was most abundant, contributing 16% of all reads in the dataset. Streptococcus_338 was most prevalent and was found to some degree in every sample.

### Baseline microbiome

DNA was successfully extracted, PCR amplified and amplicons sequenced for 411 samples, yielding a total of 956,269 high-quality reads after quality control (see Figure E1 in the online data supplement). We chose a randomly re-sampled threshold of 451 reads to ensure no bias between sample comparisons. The number of reads used discriminates very well between samples and individuals and rarefaction curves reach an asymptote indicating sufficient sampling (supplementary methods).

For the cross-sectional baseline data set, phylogenetic analyses showed the presence of 352 operational taxonomic units (OTUs), 150 of which were present in more than one subject with 21 being present at an overall abundance > 0·5%. Haemophilus_542 was the most abundant OTU overall, followed by Pseudomonas_aeruginosa_915 and Streptococcus_338 (Figure 1B). Using phylogenetic analysis we were able to confirm the Haemophilus_542 OTU to represent *H. influenzae* (see Figure E2 in the online data supplement). This approach was also attempted for Streptococcus_338, but the resulting phylogenetic trees were unable to discriminate Streptococcal OTUs at the species level (data not shown).

The most common organisms detected in sputum by clinical culture, for subjects for whom we also had 16S rRNA gene sequence data, were *Pseudomonas aeruginosa* (45·3% of subjects), *Staphylococcus aureus* (21·3 %) and *Haemophilus influenzae* (14·7%).

We compared 16S rRNA gene sequences with microbial culture by classifying OTUs as either present or absent in a subject. Comparison with DNA sequences and culture, with sequence as the putative gold standard, suggested that the calculated accuracy of cultures for *P. aeruginosa* was 71% and for *H. influenzae* was 62%, although the sensitivities were only 52% and 18%, with disagreement commonly observed in culture negative, 16S rRNA gene positive samples (Table 2). The apparent false discovery rate relative to 16S rRNA gene sequencing for culturing *P. aeruginosa* was 11%, and for S. *aureus* was 64%. There was poor accuracy (9%) and sensitivity (4%) for Streptococcal OTUs compared with culture.

**Table 2.** Comparison of rRNA gene sequences and microbial culture

| Culture ID | OTU ID | Both +ve | Both -ve | Culture -/OTU + | Culture +/OTU - | Accuracy | False Discovery Rate | Sensitivity |
| --- | --- | --- | --- | --- | --- | --- | --- | --- |
| *Pseudomonas aeruginosa* | Pseudomonas_915 | 106 (28%) | 164 (43%) | 97 (26%) | 13 (3%) | 71% | 11% | 52% |
| *Haemophilus influenzae* | Haemophilus_542 | 31 (8%) | 203 (53%) | 146 (38%) | 0 (0%) | 62% | 0% | 18% |
| *Moraxella catarrhalis* | Moraxella_945 | 16 (4%) | 313 (82%) | 51 (13%) | 0 (0%) | 87% | 0% | 24% |
| *Staphylococcus aureus* | Staphylococcus_300 | 20 (5%) | 313 (78%) | 32 (8%) | 35 (8%) | 83% | 64% | 38% |
| *Stenotrophomonas maltophilia* | Stenotrophomonas_401 | 11 (3%) | 323 (85%) | 46 (12%) | 0 (0%) | 88% | 0% | 19% |
| *Proteus vulgaris* | Proteus_1088 | 1 (0.3%) | 370 (97%) | 8 (2%) | 1 (0.3%) | 98% | 50% | 11% |
| *Streptococcus pneumoniae* | Streptococcus_338 | 13 (3%) | 23 (6%) | 343 (90%) | 1 (0.3%) | 9% | 7% | 4% |

Alpha-diversity by all measures (Shannon Diversity Index, Inverse Simpsons Index, species richness and evenness) was significantly lower if the subject was receiving prophylactic antibiotics, or if any organism had been isolated from the sample, or if mucoid *P. aeruginosa* had been isolated (Figure 2A). The patients’ gender, treatment with steroids, age, FEV1, FVC, BMI, and the years since first *P. aeruginosa* isolate were not associated with diversity.

**Figure 2.**
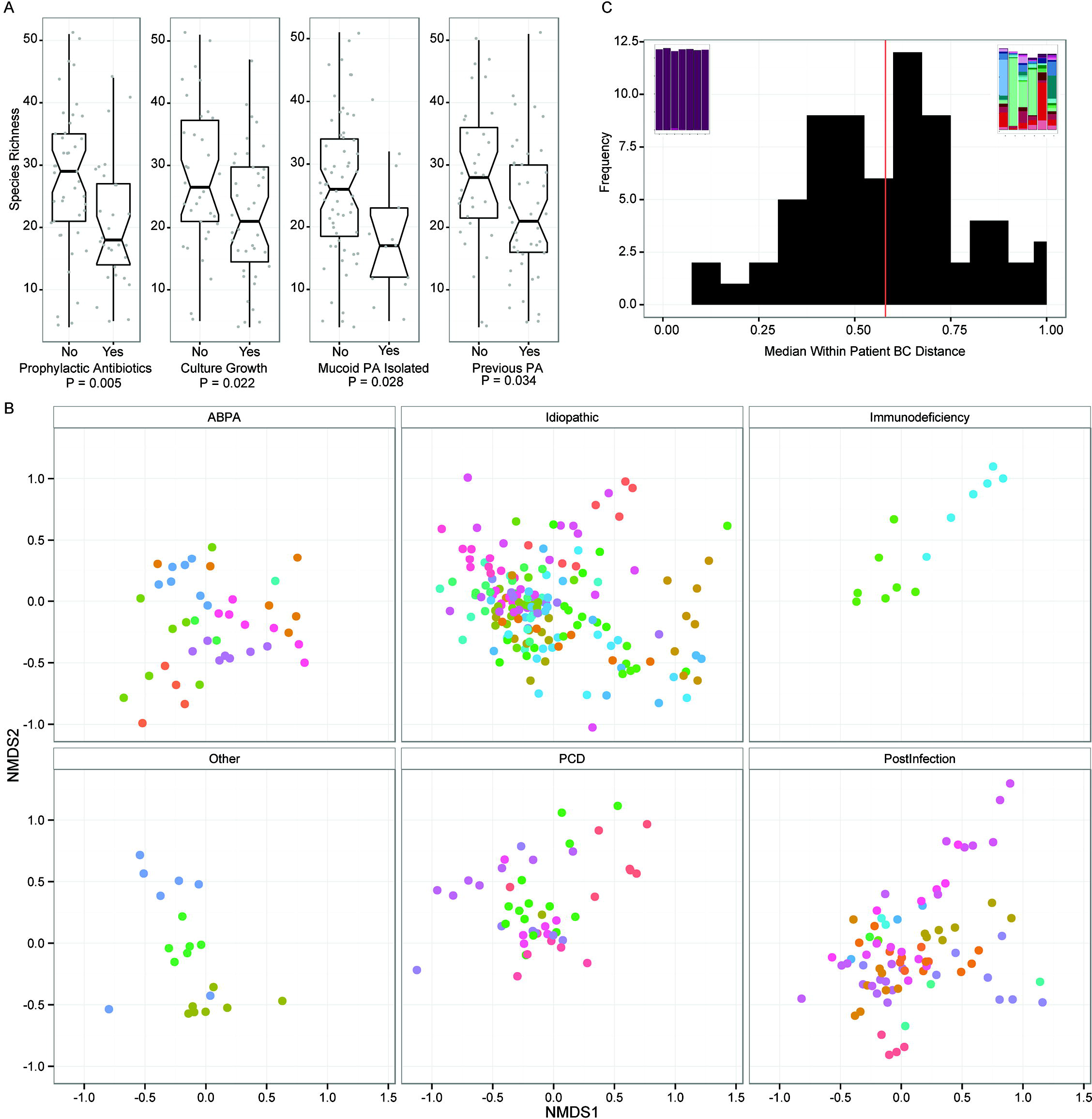
**2A.**Boxplots of species richness for cross-sectional baseline samples comparing clinical categories. Notches indicate 95% confidence interval. P values were calculated using Welch’s T test. **2B.**Non-metric multi-dimensional scaling plot of Bray-Curtis dissimilarity. This ordination plot visually represents the Adonis results. The plot has been split by underlying cause of non-CF bronchiectasis to reduce over-plotting and to enable clearer visualisation of clustering of points, although each panel can be considered to be directly overlaid upon one another. Each point represents a sample and the larger the distance between points the larger the difference in community structure of those samples. Samples from the same patient have the same colour. Samples from the same patient tend to cluster together, illustrating the high individuality. There is some separation of points evident in the underlying diseases, e.g. Post-infectious samples tend to be present in the bottom right of the plot, PCD top right, ABPA central bottom and idiopathic more widely distributed. **2C.**Histogram of the median per patient Bray Curtis dissimilarity. Bray Curtis dissimilarity was calculated for each patient with more than 3 samples and ranged from 0·12 to 0·98. The embedded stacked bar plots illustrate the patients at the two extremes, least diverse and most stable to most diverse and variable.

### βdiversity

Bray-Curtis dissimilarity was calculated in order to compare the relationship between communities of baseline samples from each individual and clinical factors We constructed permutational multi-variate ANOVA (Adonis) models of the baseline cross-sectional dataset to test the effect of individual variables on between-sample β-diversity. Variables were removed from the model if they were no longer significant, optimising the fewest number of variables that together explained the highest proportion of variance. The variance of diversity was significantly related to treatment with prophylactic Colistin (R^2^ 0·04, P=0·008) and by isolation of mucoid *P. aeruginosa* (R^2^ 0·14, P<0·001), *H. influenzae* (R^2^ 0·07, P<0·001) and *S.aureus* (R^2^ 0·04, P=0·013). These variables together accounted for 29% of the total variance in the community structure.

### Longitudinal analysis

There were 122 infective exacerbations recorded by the 64 patients that completed follow up over the 6 month period. Forty one patients had two or more exacerbations over 6 months, with 14 patients having no exacerbations and 9 patients having only one.

There was no significant difference in the exacerbation rate in patients on prophylactic antibiotics or patients with or without *P. aeruginosa*. In total, 37 exacerbations coincided with clinic visit. 18 (50%) of these were not accompanied by the growth of bacteria in culture, despite non-usage of antibiotics during the previous 30 days. Only 4/37 (11%) of the exacerbation samples were associated with isolation of a bacterium not seen in prior samples.

We used 16S rRNA gene quantitative PCR to measure the total bacterial load in the samples. The median copy number was 2·2 ×10^8^ per ml of sputum at baseline (IQR 4·8 × 10^7^ – 8·6 × 10^8^). We found no significant difference in bacterial load between the baseline, exacerbation, treatment or recovery samples (see Figure E3 in the online data supplement).

There was no significant difference in any diversity measure between exacerbation samples and paired baseline (n=13), treatment (n=5) or recovery samples (n=21) from the month immediately prior to the exacerbation (see Figure E4 in the online data supplement). There was also no difference in any diversity measure between exacerbation samples and those immediately following recovery.

A multivariate ANOVA showed that 5 9·6% of the total variation in longitudinal (β-diversity could be explained by the subject the sample was from, compared to 5·8% for the underlying disease (Figure 2B).

We calculated a per subject median Bray Curtis dissimilarity for those with 3 or more samples to give an individual range of diversity in different samples from individual subjects. A high value indicated that the microbial communities changed from month to month in relative abundance and membership, and a low value indicated that samples from the same subject were similar. The median dissimilarity was selected in order to reduce sensitivity to outlying data points and all possible pairs of dissimilarities were included. We observed a wide range of stability for the normally distributed metric (range 0·12 to 0·98; mean=0·58, median= 0·60) (Figure 2C).

The stability metric did not correlate significantly with clinical characteristics including BMI, prophylactic antibiotic treatment, number of exacerbations, average lung function (FEV1pp, FVCpp), underlying cause, and carriage of *P. aeruginosa*. Additionally there was no difference in this metric between patients who did or did not change clinically during the study.

Given the high individuality of the microbial communities, we produced per subject plots of the OTU relative abundances alongside the clinical data and quantitative PCR of the 16S rRNA gene as a proxy for bacterial load. A wide spectrum of community compositions, stability, and pathogen load were observed, (Figures 3A to 3D for examples and Figure E5 in the online data supplement for all other subjects).

**Figure 3.**
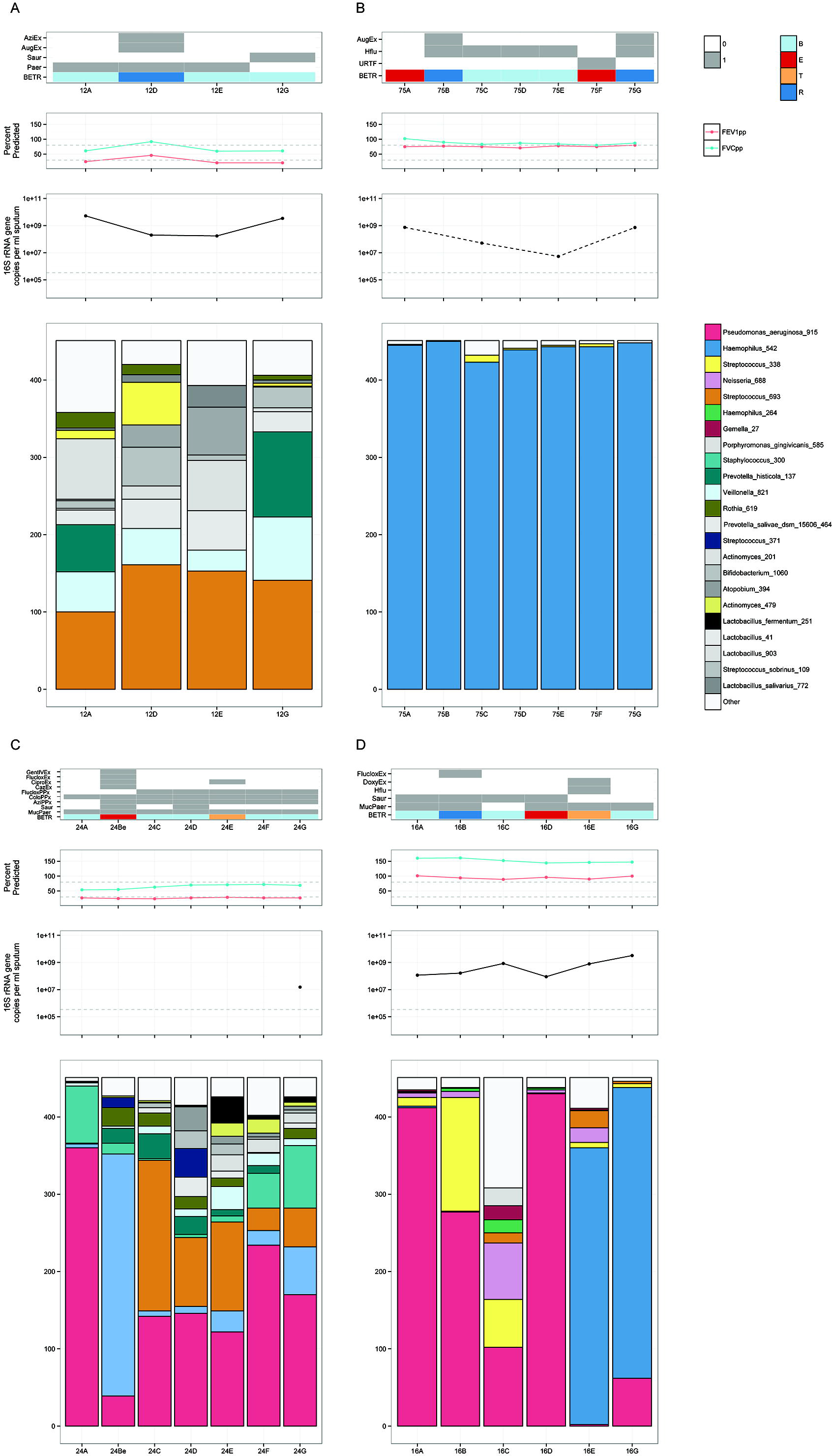
Selected subject plots. Each subject is represented by four plots, from top to bottom: clinical variables including antibiotic treatment, growth of microorganisms on clinical culture and B,E,T,R category; Lung function as FEV1 % predicted (red), FVC % predicted (green) with 30% and 80% represented by the grey dotted line; bacterial load as measured by 16S rRNA gene qPCR in copies per ml of sputum with the detection limit of the assay indicated by the grey dotted line; stacked barplots of the OTUs present in each sample. Colour coding for top 26 OTUs consistent between plots, with greyscale used for the remaining OTUs. Rare OTUs in each plot are summed as “Other”. **3A.**Subject 12: 68 yr old male with ABPA, normal BMI and 2 exacerbations during the study period. The patient had the highest median Bray-Curtis dissimilarity. Streptococcus_693 was the most abundant OTU in every sample (although not dominant) but other OTUs changed in relative abundance from sample to sample. Bacterial load changed substantially over the sampling period. **3B.**Subject 75: 64 yr old female, post-infectious, underweight and 2 exacerbations during the study period. The patient had the lowest median Bray-Curtis dissimilarity and most stable microbial community, dominated by Haemophilus_542, despite two clinical exacerbations and treatment with Augmentin. Bacterial load varied by two orders of magnitude from 10^7^ to 10^9^ copies per ml of sputum. **3C.**Subject 24: 60 yr old male, unknown BX cause, normal BMI and 2 exacerbations during the study period. The patient did show changes in bacterial community that coincided with clinical states, such as an exacerbation at time point Be associated with a large increase in abundance of Stenotrophomonas_401. Antibiotic treatment resolved the exacerbation and Stenotrophomonas_401 proportions returned to lower levels. **3D.**Subject 16: 65 yr old male with ABPA, normal BMI and 2 exacerbations during the study period. The patient had an exacerbation at samples D and E, with Pseudomonas_aeruginosa_915 initially dominant being replaced by Haemophilus_542. The proportion of Haemophilus_542 and bacterial load in the samples increased, suggesting active growth of Haemophilus_542 that was supported by coincident clinical culture of *H. influenzae*.

## Discussion and Conclusion

The study shows substantial complexity in the airway microbiome in patients with chronic bronchial infection, with frequent mixed infections and potentially important discrepancies between DNA sequencing results and classical clinical culture. The structure of microbial communities within patients was highly individual, relating only weakly to underlying disease, and often stable over the six months of the study despite the use of antibiotics and changes in clinical state.

The differences between culture and 16S rRNA gene sequencing show that the common complexity of pathogen growth is captured incompletely by standard microbial culture. In particular, the presence of *H. influenzae* appears under-recognised by culture, and *Pseudomonas spp*. and *S. aureus* were at times present in culture and not detected by sequencing. This may reflect the capacity of culture to isolate pathogens when they are present in very low numbers, and the ability of *Pseudomonas* spp. and *S. aureus* to outgrow other organisms in culture. Since the discrepancy in *S. aureus* was most marked, primer sequences were checked for specificity for this group of organisms and found to have 100% identity to the *S. aureus* target. Inefficient DNA extraction can be of concern with Gram positive organisms, although the bead-beating approach has been employed widely and in our hands efficiently lyses Mycobacteria and fungi. It has also been validated for endospore extraction, so we do not believe that inefficient DNA extraction has occurred here. It is possible however that relatively inefficient amplification of 16S rRNA gene sequences from genomic DNA of *S. aureus* against a mixed template background may have occurred.

A limitation of 16S rRNA sequencing is that particular OTUs may not define bacterial species, best exemplified by the inability to identify *Streptococcus pneumoniae* or S. *mitis* among the streptococcal OTUs. It is likely that the discrepancy between culture and the 16S rRNA gene sequencing is caused by the summing of multiple different Streptococcal species, including common respiratory commensals, into a single OTU. OTU analysis also cannot be used to define pathogenicity, and does not give information about antibiotic susceptibility.

The microbiome in our patients may reflect their advanced disease and treatment with multiple courses of antibiotics over many years. Although the literature does not yet provide a clear meta-analysis of the normal airway microbiome, the distribution and diversity of OTUs in these patients seems to differ markedly from that seen in normal subjects or those with asthma or COPD, where Gram negative anaerobes such as *Prevotella* or *Veilonella* spp. may make up to ~30% of OTU abundance (Hilty et al. 2010; Molyneaux et al. 2013; Erb-Downward et al. 2011).

It is unclear whether the lower abundance of these organisms in our patients is a result of the presence of more abundant organisms, competition with other organisms such as *P. aeruginosa* and *H. influenzae,* or whether it follows selection by treatment with antibiotics. It is possible that depletion of the commensal community may itself facilitate early pathogen introduction to community and dominance. In chronic obstructive pulmonary disease, acquisition of a new bacterial strain has been associated with exacerbations (Sethi et al. 2002). Here we lack strain level resolution, although new OTUs could only rarely be seen associating temporally with exacerbations. For example, Subject 15 Pseudomonas_aeruginosa_915 takes over from Streptococcus_338 at exacerbation, in subject 24 Stenotrophomonas_401 from Pseudomonas_aeruginosa_915 whilst there is an increase in Stpahylococcus_300 in patient 67.

Our observation that prophylactic antibiotics correlated with reduced alpha-diversity supports a community level impact of prophylactic therapy. There was however no correlation of alpha or beta diversity or bacterial load with important clinical parameters such as severity or duration of disease, suggesting that analyses of diversity are not simply a reflection of disease severity and that they should be used alongside the recognition and enumeration of pathogens.

The culture of mucoid *P. aeruginosa* was associated with changes in community structure, which might be due to the impact on the local environment in the lung of this phenotype or could be a marker of longer colonisation as mucoidy is associated with chronic colonisation (Levy et al. 2008).

Our finding that the bacterial community relates poorly to clinical state supports the results of longitudinal studies of patients with Cystic Fibrosis (Zhao et al. 2012; Stressmann et al. 2011; Tunney et al. 2013; Carmody et al. 2013; Carmody et al. 2015). Overall in these studies no, or poor associations with clinical state are seen when taking each cohort as a whole. However, as we have also demonstrated here, in subsets of patients associations can be seen. CF is a much more clinically defined disease than non-CF bronchiectasis, though strong individuality in the microbiota may also mask the influence of the microbiota on clinical state.

The present study included a number of different underlying etiologies of non-CF bronchiectasis in order to allow comparison of these and to assess whether the microbial communities supported these clinical classifications. After taking into account the strong per subject differences in microbial communities, a small proportion of the variance could be ascribed to etiology. This might indicate that in larger studies stratification by etiology would reduce the variance and increase power to detect disease-specific effects.

Here we find that *Haemophilus influenzae* and *Pseudomonas aeruginosa* are the two most common and dominant pathogens by 16S rRNA gene sequencing. In our PERMANOVA model isolation of mucoid *Pseudomonas aeruginosa* or *Haemophilus influenzae* had the greatest influence on community structure as a whole. Changes in the microbiota composition have also been demonstrated after prophylactic treatment with erythromycin in a more homogenous and milder group of bronchiectasis patients, though only when considering individuals dominated by *Haemophilus influenzae* (Rogers et al. 2014). This suggests that although underlying etiology can be important, the dominant organism present, irrespective of etiology, has greater influence and could be targeted accordingly.

Discerning the impact of individual antibiotics was difficult in this dataset, given the extremely individual microbiota and the range of different antibiotics used for prophylaxis, exacerbation, and non-respiratory reasons. Colistin was the only antibiotic that had a significant impact on community structure. It was the only nebulised antibiotic to be used frequently within the cohort, so the impact of nebulisation, where a higher concentration of antibiotic is expected local to the respiratory tract, is difficult to separate from the actions of Colistin itself. The influence of Colistin on community structure is not independent of the influence of mucoid *P. aeruginosa* as this antibiotic would be used to target the organism (Haworth et al. 2014).

We separated antibiotics into those given prophylactically for respiratory reasons, respiratory exacerbation antibiotics and non-respiratory antibiotics. Only a small number of non-respiratory antibiotics were prescribed to study patients and as these are at a lower dose and likely to have less influence on the microbial communities in the airways, these were not included in the BETR definition.

Possible explanations for the poor correlation between the sputum microbiome and clinical course include that the disease is driven by mucosal events that are poorly reflected in sputum, or that the activity of the microbiome is changing independently of bacterial load (for example through expression of virulence factors), or that exacerbations are being driven by virus or fungal infections rather than bacteria. Our study was not designed to examine exacerbation specifically, as its aim was to observe changes of the microbial community in a diverse non-CF bronchiectasis cohort over time. Consequently, only a relatively small number of exacerbation (E) or treatment (T, exacerbation with current antibiotic treatment) samples were obtained.

We were unable to obtain a full set of samples analysed by all methodologies from every individual. As can be seen in the consort diagram (Supplementary Figure E1) this was due to subjects withdrawing part way through the study, subjects being unable to expectorate sputum on a particular occasion and insufficient material for analysis at the further methodological stages of 16S rRNA gene sequencing. Lack of sputum expectoration was also an issue with T (treatment) samples as treatment reduced sputum volume. As 16S rRNA gene qPCR was applied later than 16S analyses it was only performed for subjects with sufficient sample remaining.

Sputum samples in some individuals revealed the same community every month for six months, showing consistency of community structure. The variability in other patients might result from a respiratory tract rendered inhomogeneous by advanced disease, which may confound a whole airway sample such as sputum (Jorth et al. 2015; Erb-Downward et al. 2011). More frequent sampling could establish whether changes in the microbiota not coincident with clinical change were due to sampling variability, community variability as a result of drivers with no clinical impact (e.g. competition between microorganisms) or clinical changes that occurred between visits and were not captured. Longer and more frequent sampling may also allow the development and application of more sensitive statistical and ecological methods that will also be of benefit. It is important nevertheless that the number of patients and the period of time studied here is substantial compared to previous studies of BX and other chronic suppurative lung diseases.

It is possible that serial sampling of the sputum microbiome with nucleic acid sequencing may be used to better therapeutic outcomes for patients with chronic bronchial infection. Given the extremely high individuality, longitudinal assessments of the individual patient)s microbiota during periods of health may become the best control for managing their exacerbations.

Diverse bacterial communities can be resistant to pathogen invasion, so prophylaxis with more targeted antibiotics that maintain diversity might be beneficial. Our results suggest that accurate profiling of the respiratory microbiome will lead to improvements in the understanding of the role of prophylactic antibiotics on community diversity, accurate recognition of pathogens and their interactions in complex communities, and better identification and treatment of true infective exacerbations.

Conducting even larger studies that will allow stratification of patients by underlying etiology, dominant pathogen and antibiotic treatment will increase power significantly and may lead to identification of stronger links between clinical state and the microbiota present. A number of inhaled antibiotics are in development for bronchiectasis and a better understanding of their benefits and the consequences of their use on the microbial community is needed.

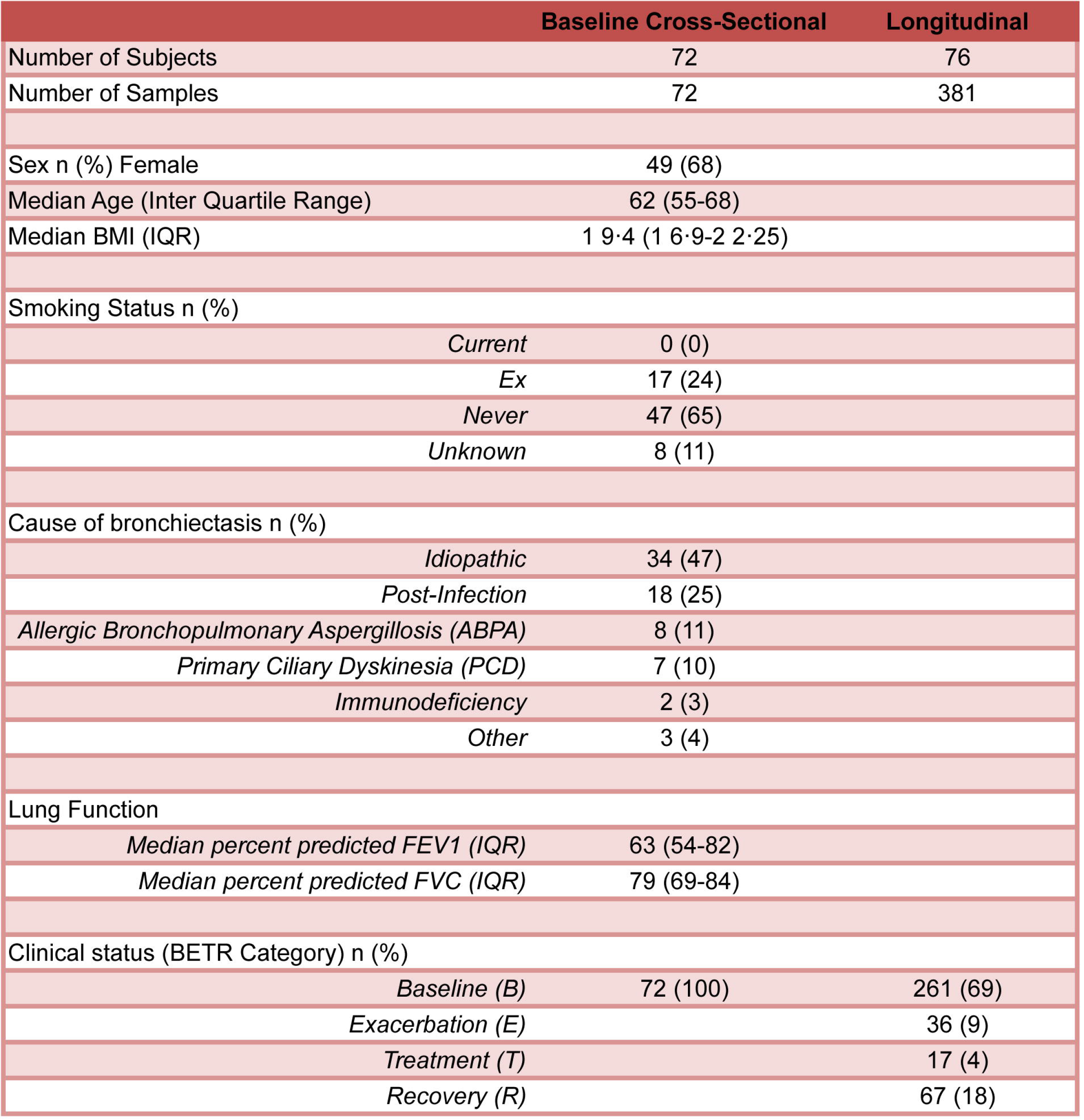

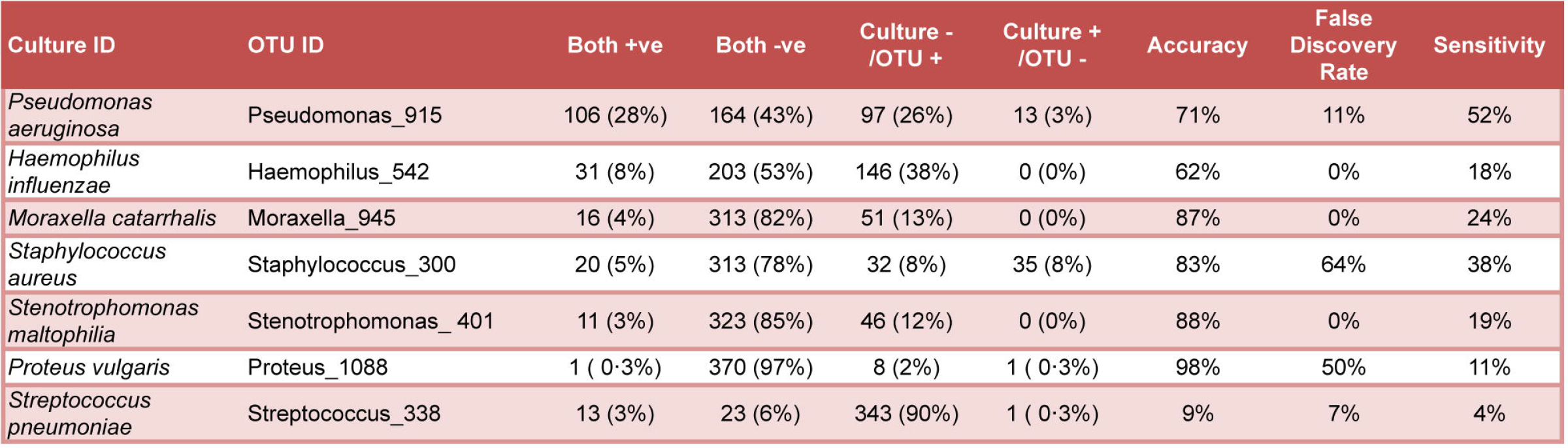

## Acknowledgements

We are grateful to the patients and staff of the Royal Brompton and Harefield NHS Foundation Trust for their assistance.

## Author contributions

MJC, ML, WOCC and MFM planned the study, based on a clinical design by ML. CH, and ML recruited patients, sampled and gathered clinical data. MJC, GKM and ET performed 16S rRNA gene laboratory studies. MJC led statistical analyses of the data with contributions from WOCC, PLJ, AJ and MC. MJC and ML wrote the first draft of the paper. All authors contributed to the interpretation of the results and the writing of the paper.

